# Multi-omics Visualization Platform: An extensible Galaxy plug-in for multi-omics data visualization and exploration

**DOI:** 10.1101/842856

**Authors:** Thomas McGowan, James E. Johnson, Praveen Kumar, Ray Sajulga, Subina Mehta, Pratik D. Jagtap, Timothy J. Griffin

## Abstract

**Background:** Proteogenomics integrates genomics, transcriptomics and mass spectrometry (MS)-based proteomics data to identify novel protein sequences arising from gene and transcript sequence variants. Proteogenomic data analysis requires integration of disparate ‘omic software tools, as well as customized tools to view and interpret results. The flexible Galaxy platform has proven valuable for proteogenomic data analysis. Here, we describe a novel Multi-omics Visualization Platform (MVP) for organizing, visualizing and exploring proteogenomic results, adding a critically needed tool for data exploration and interpretation.

**Findings:** MVP is built as an HTML Galaxy plugin, primarily based on JavaScript. Via the Galaxy API, MVP uses SQLite databases as input -- a custom datatype (mzSQLite) containing MS-based peptide identification information, a variant annotation table, and a coding sequence table. Users can interactively filter identified peptides based on sequence and data quality metrics, view annotated peptide MS data, and visualize protein-level information, along with genomic coordinates. Peptides that pass the user-defined thresholds can be sent back to Galaxy via the API for further analysis; processed data and visualizations can also be saved and shared. MVP leverages the Integrated Genomics Viewer JavaScript (IGVjs) framework, enabling interactive visualization of peptides and corresponding transcript and genomic coding information within the MVP interface.

**Conclusions:** MVP provides a powerful, extensible platform for automated, interactive visualization of proteogenomic results within the Galaxy environment, adding a unique and critically needed tool for empowering exploration and interpretation of results. The platform is extensible, providing a basis for further development of new functionalities for proteogenomic data visualization.

## Findings

Proteogenomics has emerged as a powerful approach to characterizing expressed protein products within a wide-variety of studies[1–5]. Proteogenomics, a multi-omic approach, involves the integration of genomic and/or transcriptomic data with mass spectrometry (MS)-based proteomics data. Typically, a proteogenomics-based study starts with a sample (e.g. cells grown in culture, tissue sample etc.) which are analyzed using both next generation sequencing technologies (usually RNA-Seq) and MS-based proteomics. Once assembled from RNA-Seq data, the transcriptome sequence is translated in-silico to generate a database of potentially expressed proteins encoded by the RNA. This protein sequence database contains both proteins of known sequences contained in reference databases, as well as novel protein sequences which are derived from the transcriptome sequence via comparison to reference genome sequence. These novel sequences may include variants arising from single-amino acid substitutions, short insertions/deletions, RNA processing events (truncations, splice variants) or even translation from unexpected genomic regions[2].

Parallel to the RNA-Seq analysis, tandem mass spectrometry (MS/MS) data is collected from the same sample by fragmenting peptides derived from proteolytic digestion extracted proteins. Each MS/MS spectrum contains sequence-specific information on detected peptides. Sequence database searching software[6] is used to match MS/MS spectra to peptide sequences within the RNA-Seq derived protein sequence database, providing direct evidence of expression of not only reference protein sequences, but also novel sequences. Proteogenomics provides a powerful approach to collect direct evidence of expression of novel protein sequences specific to a sample of interest, which may not necessarily be present in reference sequence databases. The value of proteogenomics has been shown in studies of cancer and disease[3–5] as well as a means to annotate genomes[7].

As with other multi-omic approaches, proteogenomics presents some unique informatics challenges[8]. For one, data from different ‘omic technologies (e.g. RNA-Seq and MS-based proteomics) must be processed using multiple domain-specific software. Once MS/MS spectra are matched to peptide sequences, further processing is necessary to ensure quality of the matches as well as to confirm novelty of any sequences identified which don’t match to known reference sequences. Finally, novel sequences must be further visualized and characterized, assessing confidence based on quality of supporting transcript sequence information and exploring the nature of the novel sequence when mapped to its genomic coding region[9].

Galaxy[10] has proven a highly capable platform for meeting the requirements of multi-omic informatics, including proteogenomics, as described by us and others[11–15]. Its amenability to integration of disparate software in a unified, user-friendly environment, along with a variety of useful features including complex workflow creation, provenance tracking and reproducibility, address the challenges of proteogenomics. As part of our work developing Galaxy for proteomics (Galaxy-P[16]), we have focused on putting in place a number of tools for the various steps necessary for proteogenomics -- from raw data processing and sequence database generation[9, 11, 12, 17], to tools for interpreting the potential impact of identified sequence variants[18] and mechanisms of regulation indicated by RNA-protein response[19]. Others have also contributed to this growing community of proteogenomic researchers utilizing Galaxy to address their data analysis and informatics needs[11–15].

However, despite this community-driven effort to develop Galaxy for proteogenomics, there are still a few missing pieces critical for complete analysis of this type of multi-omics data. Currently, there is a significant lack of tools that could filter the results from upstream proteogenomic workflows, enabling further exploration of novel sequences, including visualization of these sequences along with supporting transcript and genomic mapping information. Such a tool is critical to allowing researchers to gain understanding of variants identified, and select those of most interest for further study. Although stand-alone software options exist for viewing proteogenomics results[20, 21], no Galaxy-based tools are available to complement the rich suite of other proteogenomic software available within this environment. To this end, we have developed a Multi-omics Visualization Platform (MVP) which leverages Galaxy’s amenability to customized plugin tools and enables exploration, visualization and interpretation of multi-omic data underlying results generated by proteogenomics.

### Operation

MVP is built as a Galaxy visualization plugin[22], based primarily on Javascript, with HTML5 and Cascading Style Sheets (CSS) to create the interactive user interface (See Methods below for details). In Galaxy, visualization plugins require the software application (here MVP), and defined data types which act as inputs. Once data types are defined, a sniffer function in Galaxy enables the user an option to launch the plugin when appropriate data types are detected and available within the active History.

MVP operates using three separate SQLite databases for its primary input. The tool also reads the raw MS/MS peak lists (formatted as Mascot Generic File, MGF) and FASTA protein sequence database viewing results within the MVP interface. **Figure 1** shows the inputs to MVP. The SQLite databases are structured to deliver data to MVP efficiently, enabling interactive operation by users through the MVP interface. These have also been developed to present the necessary data to MVP to enable a full exploration of proteogenomics results, including evaluation of MS/MS data supporting the identification of novel peptide sequences and visualizing peptide sequences mapped to corresponding transcript and genomic coding sequences. As the primary database utilized by MVP, we have defined an mz.sqlite datatype in Galaxy which utilizes results from upstream sequence database searching software that generates output with peptide spectrum matches (PSMs), which assigns peptide sequences to each MS/MS spectrum. The mz.sqlite is generated by the mzToSQLite Galaxy tool for parsing information on: a) PSMs contained in standard mzIdentML[23] output files; b) corresponding information on MS/MS data from processed raw data files in the standard MGF format[24]; and c) the protein sequences contained in the FASTA-formatted database used for the sequence database searching.

**Figure 1.**
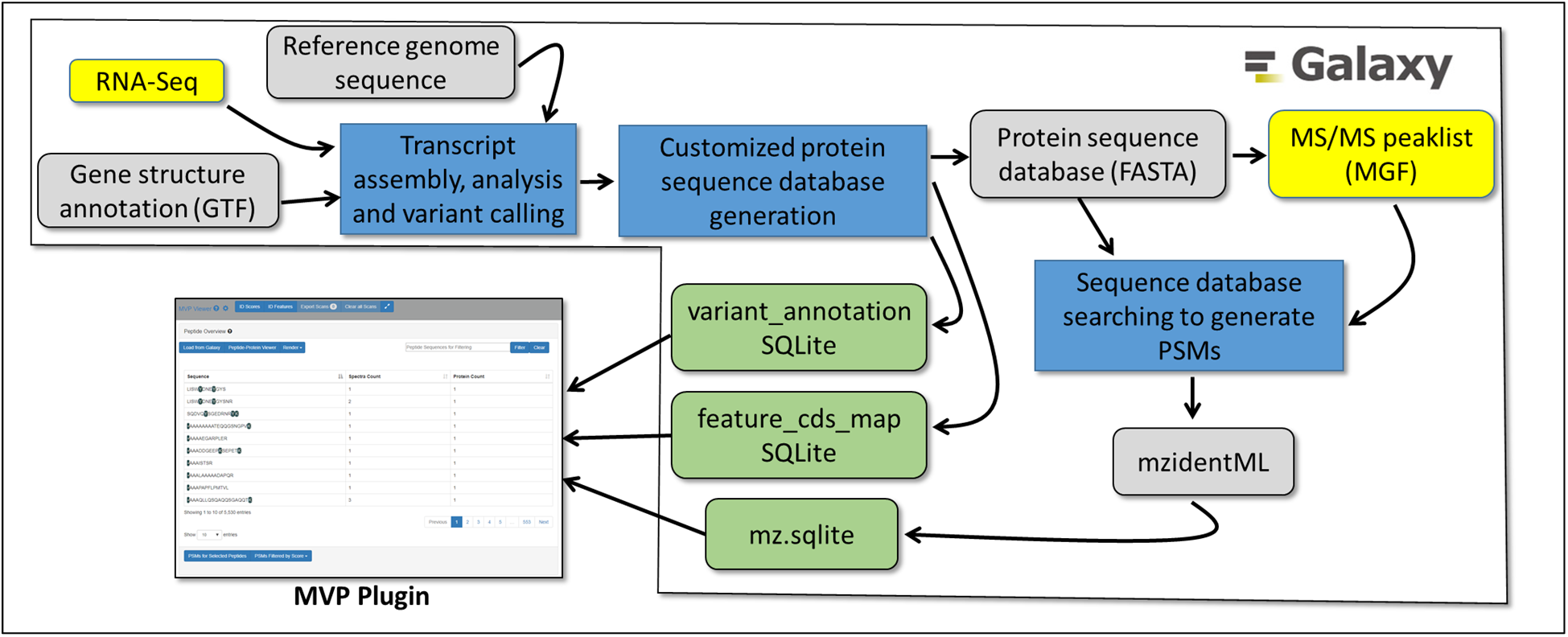
Inputs to the MVP Galaxy Plugin.

We have also developed accessible workflows and training material for generating upstream results which ultimately provide the inputs necessary for MVP operation, as shown in the workflows depicted in **Figure 1**. These include workflows for generating protein FASTA databases from RNA-Seq data, as well as matching of MS/MS data to peptide sequences via sequence database searching. Instructions for accessing these resources are described below in the Accessibility section.

Two additional SQLite tables allow MVP to display information critical for proteogenomic analysis. The variant_annotation table provides MVP information necessary to display and explore novel peptide sequences identified by matching MS/MS to the RNA-Seq derived protein sequence database. The variant_annotation table contains detailed information on how a novel peptide sequence differs from reference proteins, based ultimately on a comparison to reference genome data. The variant_annotation table is formatted with four columns: 1) name TEXT, which is the identifier of the protein with supporting PSM data, including annotation describing the nature of the novel sequence variant (single amino acid substitution, InDel etc); 2) reference Text, which is the identifier of the reference protein sequence matching protein described in column 1; 3) cigar TEXT, which is a Compact Idiosyncratic Gapped Alignment Report (CIGAR[25]) text string describing the sequence differences between the reference protein and the sequence variant. CIGAR is a standard annotation method which borrows the syntax from the sequence alignment map (SAM) format[26], but uses only the operators: =, X, I, D (equal,variant,Insertion,Deletion); 4) annotation TEXT, which provides information on the exact nature of the amino acid changes between the novel variant and the reference. **Table 1** provides an example of the structure and format of the SQLite variant annotation table.

**Table 1.** Example data structure and format for the variant_annotation table

|  |  |  |  |  |
| --- | --- | --- | --- | --- |
| Example entry | variant_annotation table columns |  |  |  |
|  | name TEXT* | reference TEXT | cigar TEXT* | annotation TEXT |
| **Single amino acid substitution** | ENSMUSP00000092502_E59G.Q267R | ENSMUSP00000092502 | 58=1X207=1X26= | E59G,Q267R |
| **Nucleotide insertion/deletion resulting in frame shift** | P_001316432_24_C>cA | P_001316432 | 7=99X | C>cA |
| **In-frame amino acid insertion** | P_001316432_24_C>cATG | P_001316432 | 7=1I99= | C>cATG |
| **In-frame amino acid deletion** | P_001316432_24_CATG>c | P_001316432 | 7=1D97= | CATG>c |
| **Structural variant (fusion)** | TAF1D_SNORA40 | TAF1D | 24=14X | chr3:18100200+,chr6:28233455- |
| *MVP reads only the name TEXT field and cigar TEXT fields for identified sequence variants. The reference TEXT and annotation TEXT fields are included to identify the reference gene and a free-form description of the nature of the annotated variant, respectively, and are not directly read by MVP.<br>For the example variants provided, the cigar TEXT string describes the following:<br><u>Single amino acid substitution:</u> For this example the cigar text depicts an amino acid (AA) sequence that contains 58 AAs that match the reference exactly, followed by 1 AA difference (X), followed by 207 AAs that match exactly, followed by 1 AA difference, followed by 26 AAs that match exactly.<br><u>Nucleotide insertion/deletion causing a frameshift:</u> For this example, the cigar text depicts an AA sequence where the first 7 AAs exactly match the reference, followed by 99 mismatched amino acids due to a single nucleotide insertion into the coding sequence which causes a change in reading frame for the remainder of the protein sequence.<br><u>In-frame amino acid insertion/deletion:</u> Two examples are shown, where three nucleotides are inserted or deleted within the coding sequence. For the Insertion, the nucleotides ATG are inserted at a position such that the first 7 AAs match the reference exactly, followed by insertion of 1 mismatched AA, followed by 99 AA that are still coded in-frame and exactly match the reference; For the Deletion example, the nucleotides ATG are deleted, removing a single amino acid. Again, the first 7 AAs match exactly, followed by a deletion of a single AA and an exact match of the remaining 97 AAs in the protein sequence.<br><u>Structural variant:</u> This example shows the depiction of a structural variant (fusion), which results from the joining of two distinct annotated genes (TAF1D and SNORA40). The cigar string depicts the nature of this fusion – the first 24 AAs match the reference TAF1D sequence exactly, with the next 14 AAs mismatching due to these being from the fused SNORA40 sequence. The annotation TEXT field provides information on the chromosomal locations of the fused genes which code for the variant protein. |  |  |  |  |

The third SQLite table is the feature_cds_map, which provides information necessary to map protein sequences with supporting PSM information to genomic coordinates. This mapping is required in order to view identified peptide sequences (variant or reference) against the genome and also corresponding transcript sequence data derived from supporting RNA-Seq data in proteogenomic studies. This table contains information specific to a genome build (e.g. hg19, hg38, mm10) specified by the user within the upstream workflow when assembling transcript sequences and generating the protein sequence database. Essentially the table feature_cds_map provides a mapping of the expressed amino acid sequence for proteins inferred from PSMs to each of the exons in the reference genome coding for the protein. Notably, MVP is amenable to any organism where a reference genome build is available, such that it is useful for a wide-variety of proteogenomics studies.

For each coding exon of a translated protein, the feature_cds_map table is formatted with these columns: 1) name TEXT, which is the identifier of the reference or variant protein with supporting PSM data; 2) chrom TEXT, which is the identifier for the reference genome chromosome coding the reference protein; 3) start INTEGER, the location of the start site for the coding region in the chromosome; 4) end INTEGER, the location of the stop site (end) for the coding region in the chromosome; 5) strand TEXT, which identifies the coding DNA strand + or - for the protein sequence; 6) cds_start INTEGER, the codon sequence at the exon start site coding the protein; 7) cds_end INTEGER, the codon sequence at the exon end site coding the protein.

**Table 2** provides an example of the structure and format of the SQLite feature_cds_map table. It should be noted that this table can also represent structural variants that are common in some cancers[27], where the variant protein maps to exons that are found on different chromosomes and/or different strands from each other. These differences would be annotated in the appropriate columns within the feature_cds_map table.

**Table 2.** Example data structure and format for the feature_cds_map table

| Table 2. Example data structure and format for the feature_cds_map table |  |  |  |  |  |  |  |
| --- | --- | --- | --- | --- | --- | --- | --- |
| Example entry | feature_cds_map table columns |  |  |  |  |  |  |
|  | name TEXT | chrom TEXT | start INTEGER | end INTEGER | strand TEXT | cds_start INTEGER | cds_end INTEGER |
| Exon* | ENSMUSP00000000033 | chr7 | 142655653 | 142655810 | - | 0 | 157 |
| Exon* | ENSMUSP00000000033 | chr7 | 142654299 | 142654448 | - | 157 | 306 |
| Exon* | ENSMUSP00000000033 | chr7 | 142653815 | 142654052 | - | 306 | 543 |
| Structural Variant (Fusion)# | TAF1D_SNORA40 | chr3 | 18100128 | 18100200 | + | 1 | 24 |
| Structural Variant (Fusion)# | TAF1D_SNORA40 | chr6 | 28233413 | 28233455 | - | 25 | 39 |
| *In this example, the reference ENSMUSP00000000033 has three exons which are mapped to chromosome 7. The table uses coding sequence (CDS) start and end locations since the codon sequence for an amino acid can be potentially split across exons. For variants such as single AA substitutions and InDels, the number of nucleotides between start and end locations may differ from the number of nucleotides in the reference CDS.<br>#An example depiction of a structural variant (fusion) is also provided. Here, two entries are included which provides information on genomic localization of the two coding sequences that were fused together to create the fusion protein, one from the positive strand of Chromosome 3 and the other from the negative strand of Chromosome 6. |  |  |  |  |  |  |  |

The MVP plugin is invoked from the mz.sqlite datatype generated within a Galaxy workflow. **Figure 2** shows the steps to invoking MVP from an mz.sqlite item within an active Galaxy History. MVP utilizes the visualizations registry and plugin framework in Galaxy[28]. The configuration and code for the plugin is placed in the visualizations plugin directory within the Galaxy installation. The configuration file for the MVP plugin defines the datatypes used as input and the code which is used to generate the interactive HTML-based interface. Once the plugin is launched, it then interacts with other input datatypes, if present, within the active History via the Galaxy API. These include the SQLite variant_annotation table and the feature_cds_map needed for characterizing variants and mapping peptides to genomic coordinates, respectively. These datatypes are automatically accessed through functions built into MVP (e.g. tools for visualizing peptides mapped to genomic sequences as described in the Functionality section below). MVP also utilizes the processed MS/MS peak lists along with the FASTA protein sequence database used for generating PSMs and contained in the active History. These inputs provide necessary data for viewing PSMs and supporting data as well as peptide data mapped against full protein sequences.

**Figure 2.**
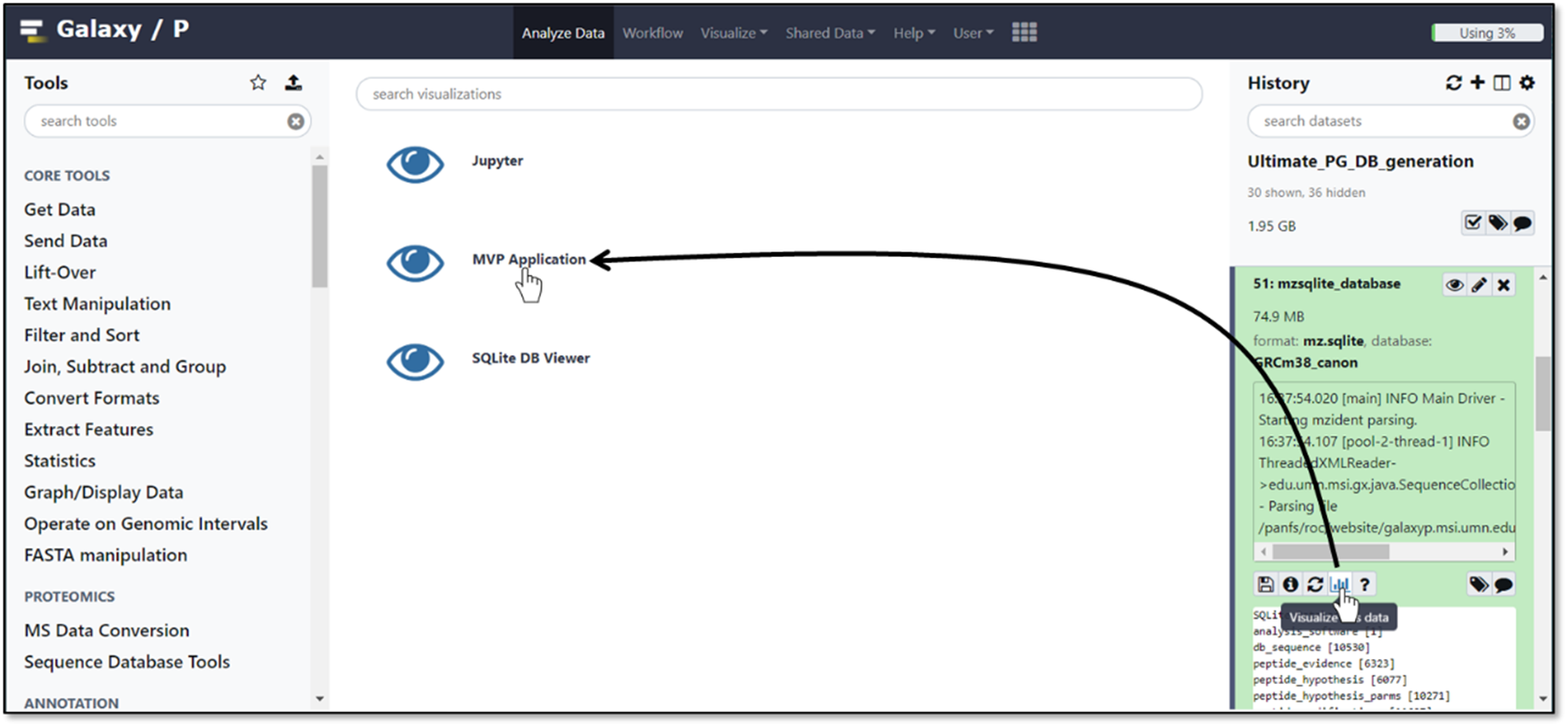
Launching MVP from an mzSQLite datatype within an active Galaxy History.

MVP makes use of two existing tools to provide functionalities critical for proteogenomics data exploration. First, it uses the JQuery-based Lorikeet viewer[29], which provides interactive viewing of annotated MS/MS spectra based on results from sequence database searching programs. Lorikeet renders a plot of peptide fragment ions and annotation from the PSM data generated from the sequence database search, offering users the ability to zoom and select or de-select specific annotation information for the peptide. This allows users to visually explore data quality for PSMs of interest, including those putatively matching novel sequences[9]. Lorikeet functionality is described in more detail below in the Functionality section.

MVP also leverages the Integrated Genomics Viewer JavaScript framework (IGVjs)[30]. Using the genomic reference sequence information contained in the feature_cds_map file corresponding to identified peptides sequences, IGVjs can be automatically launched within the MVP interface. IGVjs offers interactive viewing of peptides mapped against the reference genome, and also can add additional tracks for standard-format sequence files (e.g. BAM, ProBAM[31], BED) if present in the active Galaxy History, interacting through the Galaxy API. IGVjs provides users a flexible tool for viewing all levels of information for an identified peptide sequence -- from genomic mapping to the supporting transcript sequencing information.

It is important to note that the outputs generated by MVP processing can be used as an input for further analysis within a Galaxy history. For example, selected peptide sequences (e.g. novel sequences verified within MVP) can be sent back to the active History via the Galaxy API where they can be further processed using Galaxy tools as desired by the user. Annotated MS/MS spectra for PSMs of interest visualized within the Lorikeet viewer can also be downloaded to the desktop as a .png formatted file.

### Functionality

In order to demonstrate functionality of MVP, we have chosen a previously published dataset containing MS-based proteomic and RNA-Seq data generated from a mouse cell sample[32]. This dataset provides representative multi-omic data mimicking other contemporary proteogenomic studies, and a means to illustrate how MVP enables data exploration steps commonly pursued by researchers. The tour of MVP functionality presented here works from input data produced within a Galaxy workflow. We have made workflows available to generate input data needed for a user to explore the functionality of MVP, along with documentation describing their operation (see Accessibility section below for instructions and links to access this data).

We begin with a view of the MVP user interface, launched as a plugin from an mzSQLite data input within the active Galaxy History (See **Figure 2** above). MVP is initially launched within the center pane of the Galaxy interface, with the option of launching in a dedicated browser window. **Figure 3** shows the MVP interface initially presented to the user, where the entire set of PSMs contained in the mzSQLite database are first made available.

**Figure 3.**
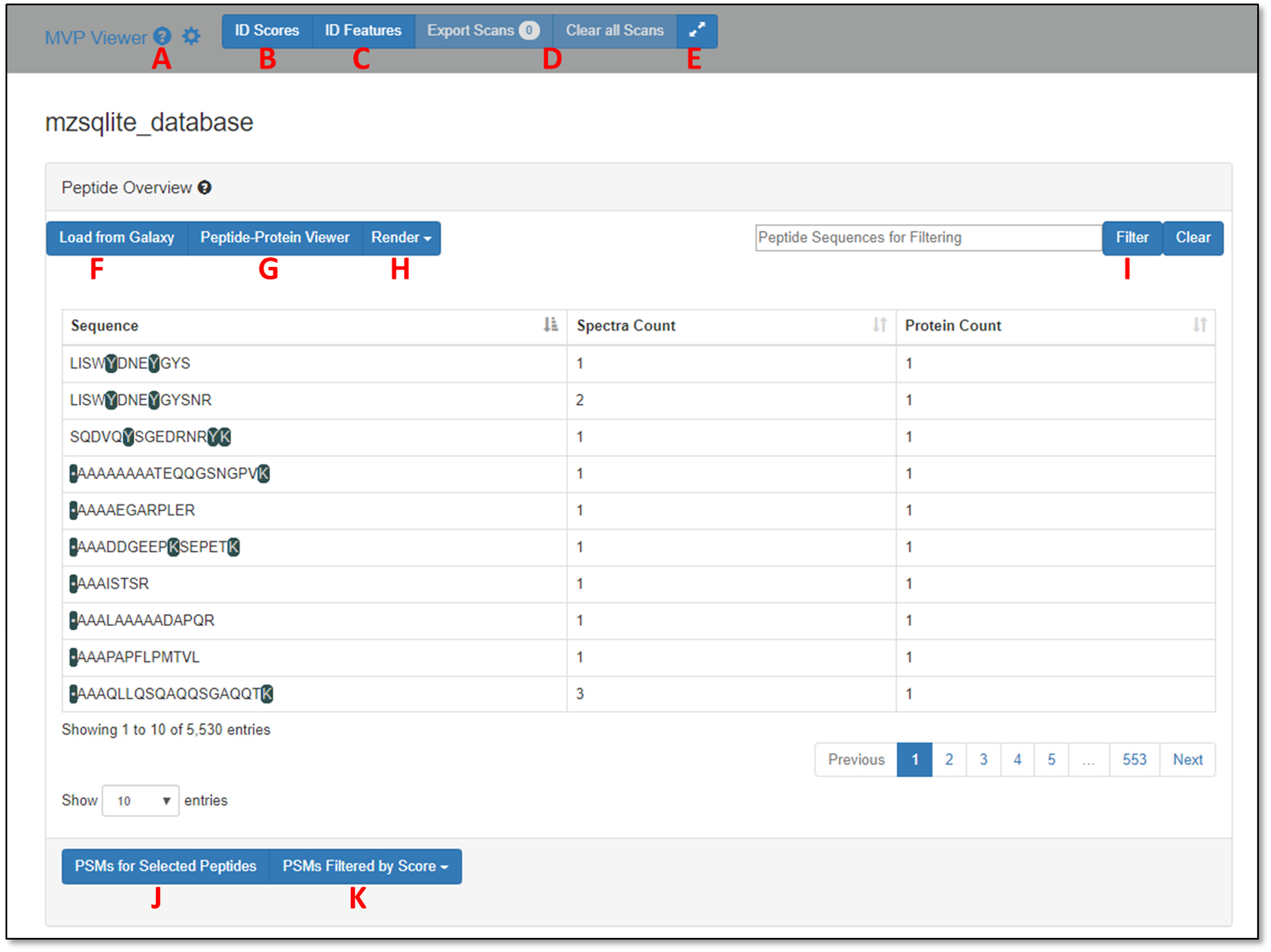
The MVP interface after initial launch from Galaxy.

The user is first presented with an unfiltered PSM-level view of all data contained in the mzSQLite data table. Peptide sequences are shown in the main viewing pane. Colored sequence elements within the peptide are those containing a modification -- in this case the sample was labeled with the iTRAQ reagent[33] on n-terminus, lysine (K) and some tyrosines (Y). Mousing over these highlighted sequence features provides a description of the nature of the modification.

From here, a number of interactive functionalities are available to the user, each labeled in **Figure 3**:

A. **“Tool Tips”** are included in each section of the software, launched by clicking any of the question mark icons. These contain a brief overview of the purpose behind the associated software feature and its functionality.
B. **ID Scores** opens a graphical description of the score distributions for PSMs passing the false discovery rate (FDR) threshold (usually set at 1% in the upstream sequence database searching algorithm generating the PSMs), and is based on information contained in the mzIdentML file outputted from the upstream sequence database searching algorithm.
C. **ID Features** provides the user a means to select the scores and data features which are displayed with each PSM, including advanced metrics (e.g. number of consecutive b or y-ions matched, total MS/MS ion current etc.) which may be useful for more advanced filtering and evaluation of quality of MS/MS matches. This is an updated embodiment of our prior description of a tool called PSM Evaluator (PSME)[9].
D. **Export Scans** provides a means to send selected PSMs in the main table back to the active Galaxy History for further analysis. **Clear Scans** deselects any selected PSMs and resets the view.
E. **The double arrows** expands and opens MVP into a new window out of the Galaxy center pane.
F. **Load from Galaxy** allows a user to import a list of peptide sequences in tabular format which have been pre-filtered and processed within the active History, for further characterization in MVP. For example, a list of pre-filtered peptides containing novel sequence variants could be imported for further analysis using this feature.
G. **Peptide-Protein Viewer** enables viewing of selected peptides aligned with their parent intact protein sequence, as well as providing a path to visualizing peptide sequences mapped to genomic sequences along with supporting transcript data (see Figure 5 and description below).
H. **Render** generates a visualization of annotated MS/MS data for all peptides shown in the current view, using the Lorikeet viewer (see Figure 4 and description below). For peptide sequences matched to multiple PSMs, the user can select which PSM is displayed in Lorikeet based on available score information. The PSM with the best score for the metric selected is then shown.
I. **Filter** allows for searching and filtering of the dataset based on input of a known sequence of interest. This will return a listing of the peptide of interest if it is contained in the mzSQLite database.
J. **PSMs for Selected Peptides** will open the annotated MS/MS in the Lorikeet viewer for any peptides selected in the center viewing pane. Multiple peptides can be selected by holding the Ctrl key and clicking each peptide of interest. For peptides with multiple PSMs, the best scoring MS/MS will be opened for viewing based on user selections from the Render button.
K. **PSMs filtered by Score** allows the user to filter either the global set of PSMs (all PSMs) or only those shown in the current screen using Boolean operators. Peptides can be filtered by a score (e.g. Confidence score from the sequence database searching program) and/or other more advanced metrics (e.g. number of concurrent b and y ions identified, total ion current etc).

**Figure 4.**
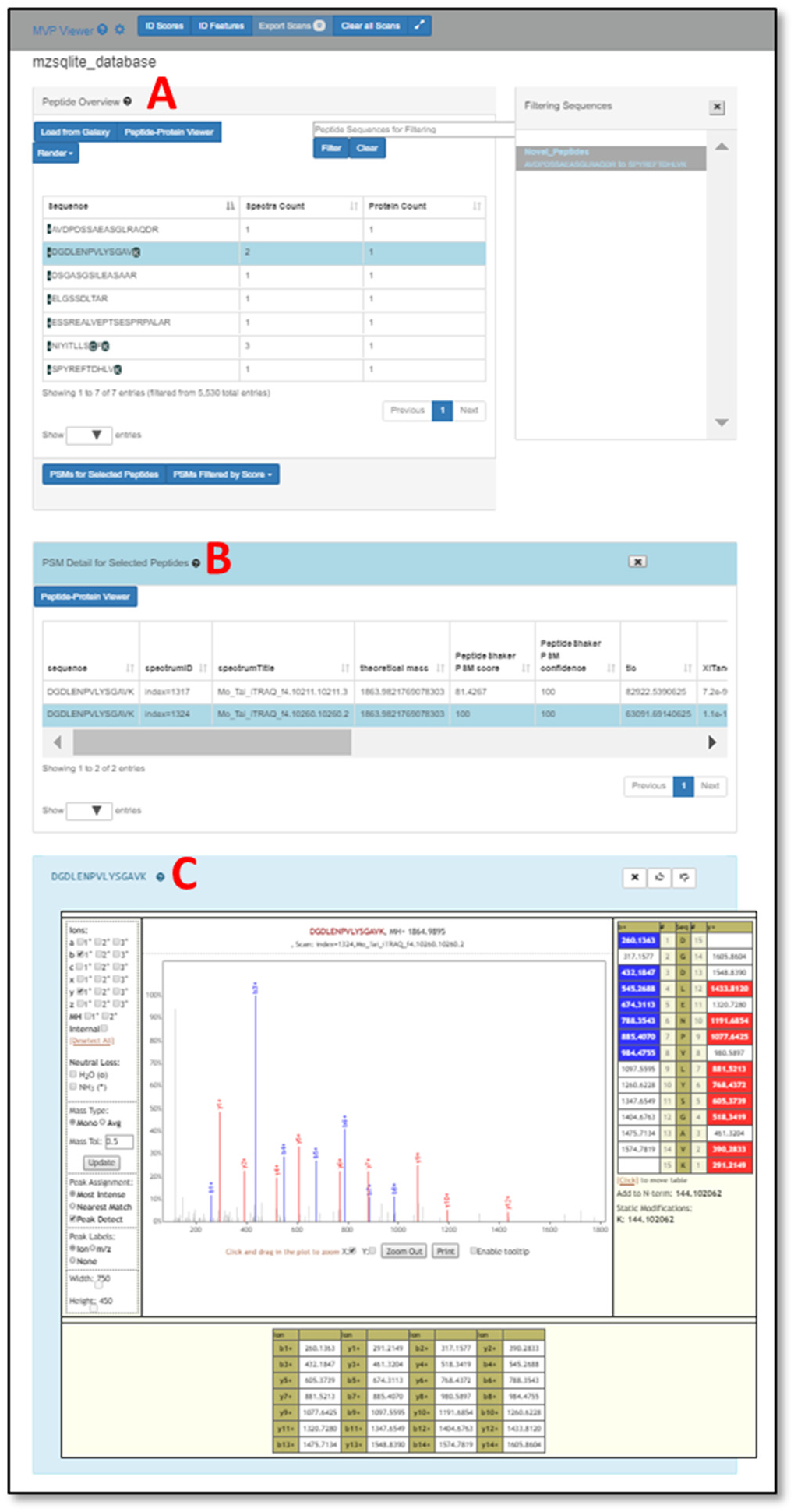
Viewing of annotated MS/MS data supporting selected PSMs

**Figure 5.**
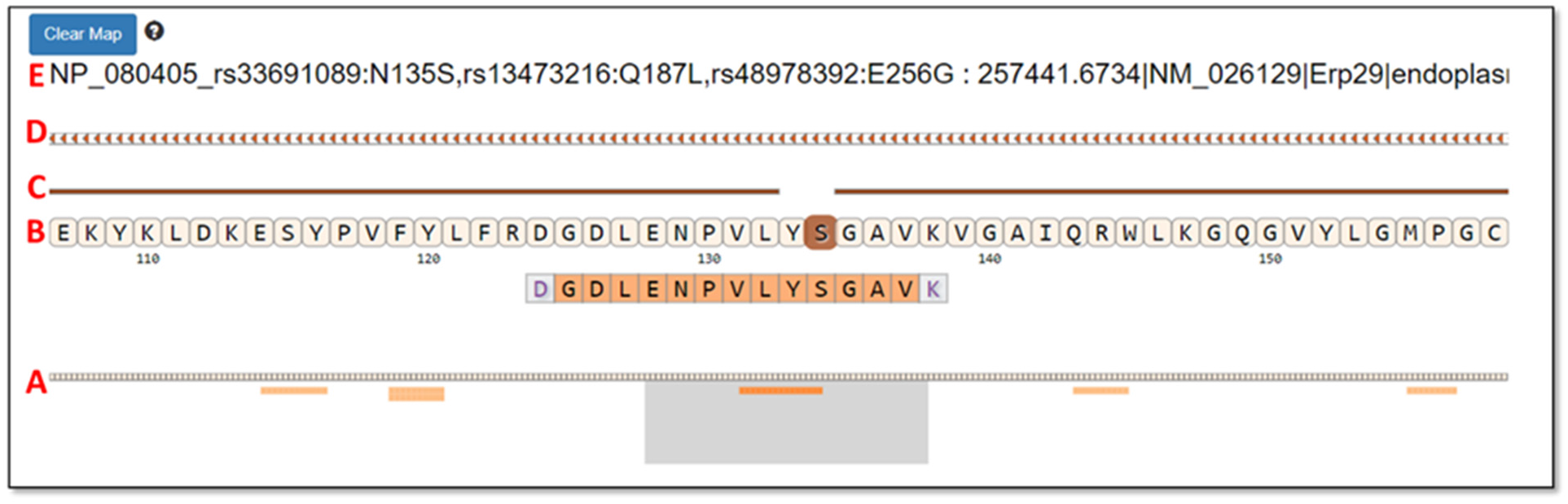
Example data shown within the Protein-Peptide viewer functionality.

To demonstrate functionalities within MVP, we follow the first step of the “Load from Galaxy” feature, loading from the active history via the Galaxy API a list of seven peptides identified in a proteogenomics workflow and confirmed as matches to variant sequences translated from variant transcript sequences. **Figure 4** displays this example data where the list of variant peptides are shown in the Peptide Overview window (Labeled A in Figure 4). One of these peptides (sequence DGDLENPVLYSGAVK) has been selected in this list, and the button “PSMs for Selected Peptides” clicked to display the two PSMs that matched to this sequence, along with associated scoring metrics (Labeled B in Figure 4). Double-clicking on one of these PSMs opens the Lorikeet MS/MS viewer (Labeled C in Figure 4). Lorikeet[29] renders MS/MS spectra providing a visualization of the annotated spectra which led to a PSM using the upstream sequence database searching software. Figure 4C shows an example PSM, where the blue and red colored m/z peak values correspond to amino acid fragments which would be predicted to derive from the peptide sequence identified by this PSM. The higher the number of peaks within the spectrum matching to predicted amino acid fragment m/z values, the higher quality and confidence of the PSM and identified peptide sequence. Lorikeet is interactive, capable of magnifying spectral regions of interest, selecting desired predicted fragment types to display, and adjustment of data parameters (such as mass accuracy of acquired data) which are commonly used in assessing PSM quality. Within MVP, this tool provides a necessary function for users to view PSMs of interest, particularly useful for assessing the accuracy of matches to variant peptide sequences in proteogenomic applications, which require extra scrutiny compared to matches to reference peptides[9].

Once the quality of a given PSM has been adequately assessed, a common user need is viewing the peptide sequence in the context of its aligned protein sequence. MVP provides this functionality, by selecting the Peptide-Protein Viewer button (available in the Peptide-Protein Viewer pane, labeled B in Figure 4). This provides a listing of all proteins within the FASTA database used for generating PSMs which contain the selected peptide sequence. For example, **Figure 5** shows the Peptide-Protein Viewer for DGDLENPVLYSGAVK (peptide sequence from Figure 4 above), along with the aligned protein sequence (the protein Erp29) containing this peptide.

Below we describe briefly the functionalities within the Peptide-Protein Viewer, following the labels (A-D) shown in Figure 5:

A. This viewing track shows all peptides within the protein sequence identified by PSMs within the dataset, depicted as lighter color lines below the aligned, complete protein sequence. The peptide sequence that was originally selected from the Peptide Overview window is depicted in a darker orange color, while other peptides also identified from the protein are in lighter orange. The gray box can be slid left or right across the entire aligned protein sequence, providing an interactive, detailed view of the peptides and aligned protein sequence contained within the box in the section labeled B in the figure.
B. For the peptide and aligned protein sequence contained in the gray box below, this section displays a zoom-in on the amino acid sequences for this particular region - both the peptide identified from a PSM and the parent protein sequence. For the example peptide, the Serine (S) is darkened within the aligned protein sequence. This indicates a position within the parent protein that differs from the reference protein, here indicating a single-amino acid substitution within the identified peptide. The n-terminal amino acid (D) and the c-terminal amino acid (K) are depicted in gray because these carried modifications due to iTRAQ reagent[33] labeling on these residues for the example dataset used for this demonstration[32].
C. The line above the protein sequence indicates alignment to the reference. If there is complete alignment, the line is solid. For any region where the protein sequence differs from the reference (based on annotation extracted from the variant_annotation table), this line is broken. In the example, the line breaks at the site of the single amino acid substitution.
D. This track in the viewer represents the genomic coding region for the protein being displayed, extracted from the feature_cds_map table. Clicking on this track will open the IGVjs browser, set to display the corresponding genomic region.
E. Here the header for the protein is shown, read from the FASTA sequence database file. For this particularly protein (Erp29), the start of the header describes positions within the sequence containing amino acid variants, including the N to S substitution detected at position 135 in the selected peptide.

From the Peptide-Protein Viewer another critical and unique functionality for proteogenomic data exploration can be launched -- specifically visualization of peptide sequences mapped to the genome and corresponding transcript sequencing data. This functionality provides MVP the capabilities to view multi-omic data, beyond the protein-level exploration of PSMs and protein sequence alignments of the Peptide-Protein Viewer. The visualization is opened automatically by clicking on the chromosome track (part D in Figure 5) which opens the IGVjs tool embedded within the MVP interface. IGV provides a rich suite of interactive functionalities, which are described in the available documentation[30]. Here we focus on several IGV features which are of most interest to proteogenomics researchers.

**Figure 6** shows the IGV viewer, with several tracks of information loaded for investigation from the active Galaxy History, investigating the genomic region coding for the peptide DGDLENPVLYSGAVK shown in Figure 5 above. This display shows information related to this peptide sequences, genomic, transcriptomic and proteomics.

A. This shows the reference sequence of the DNA coding strand for the peptide, with chromosomal position numbers shown above.
B. This track details the three-frame translation of the DNA sequence. The user can select either “forward” or “reverse” direction for translation. For the indicated peptide, the translation direction was reversed, proceeding in the direction indicated by the arrow in the figure. The frame coding the identified peptide is shaded in red.
C. Track C shows the identified peptide mapped to the genomic coordinates shown above. The arrows indicate the direction of translation against the genomic coding sequence.
D. This track summarizes the transcript sequencing reads assembled from the RNA-Seq data. This allows the user to assess the quality of supporting transcript information that led to the generation of the peptide sequence that was matched to the MS/MS data. The assembled transcript sequence read data was loaded from the active Galaxy History, contained in a standard format .bam file for assembled transcript sequencing data.

**Figure 6.**
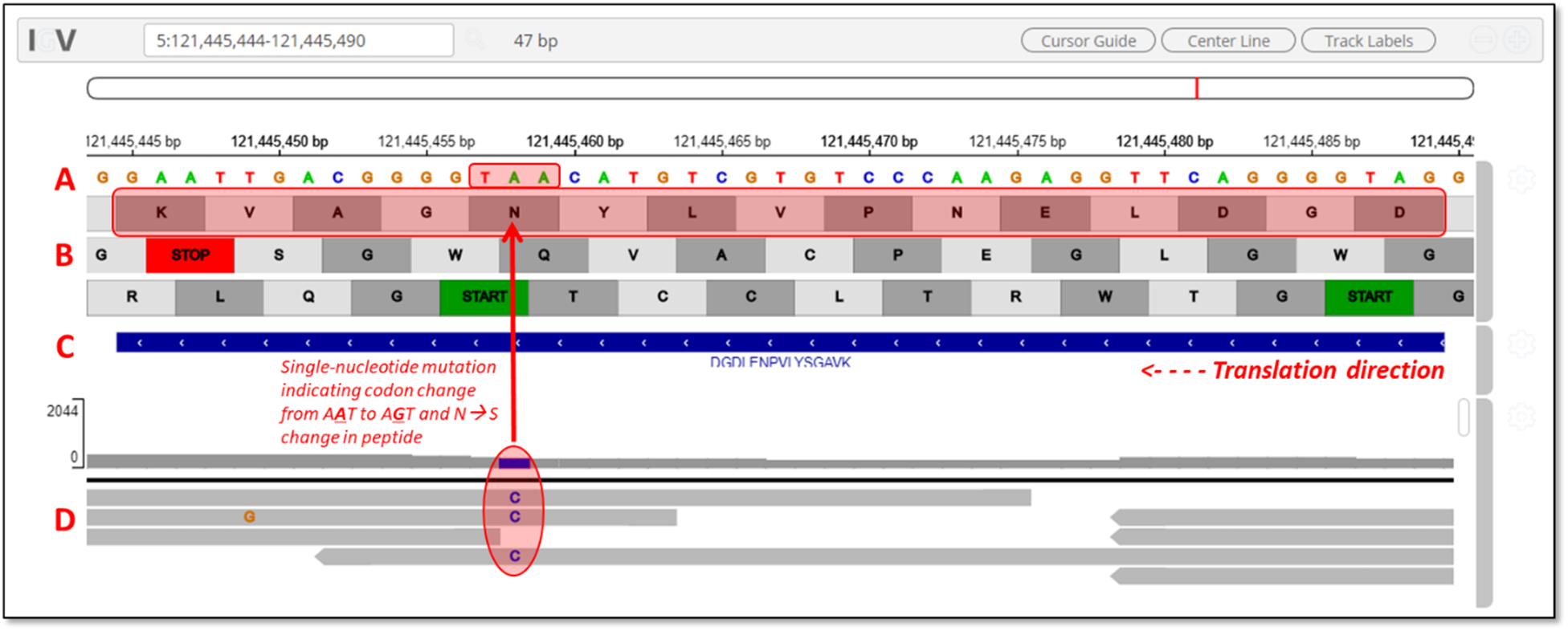
Snapshot of visualization of peptide and transcript mapping to the genome in IGV

The peptide identified here contains a single amino acid variant at the serine in position 11 from the n-terminal end of the peptide. Ordinarily, this peptide contains an asparagine at this position, as indicated by the codon A**A**T indicated in the reference DNA sequence track (Track A in Figure 6). The assembled transcript data indicates a single-nucleotide mutation within this codon, showing a C nucleotide substitution within several of the assembled reads on the negative strand (Track D in Figure 6). This substitution would indicate a complementary change at the DNA codon sequence to A**G**T, which codes for serine. The MS-based proteomics data has confirmed the expression of this variant peptide sequence. The embedded IGV tool within MVP allows users to explore this data and understand the nature and quality of the multi-omics data supporting the identification of this variant protein. **Additional File 1** shows another example of a peptide variant displayed in MVP and the IGV viewer – with this sequence containing both a single amino acid substitution and spanning a splice junction.

We have provided a guided description of the main features offered by the MVP tool. Although many powerful features are already in place to meet the requirements of proteogenomic data analysis, MVP has been developed as an extensible framework with much potential for continued enhancement and new functionalities. Tools are already implemented in Galaxy for peptide-level quantification using label-free intensity-based measurements[34, 35], which could be added to the information available for PSMs, enabling users to assess quality of abundance measurements and potentially filter for PSMs showing differential abundance across experimental conditions. The HTML5, JavaScript, and CSS-based architecture of MVP provides the ability to interact with RESTful web services offered by complementary tools and databases, as well as with the Galaxy API. We envision extending functionalities in MVP, offering users the ability to query knowledge bases[36, 37] to explore known disease-associations, interaction networks and biochemical pathways of proteins of interest. MVP also has the potential to display visualizations returned from queries against these knowledge bases. Validated peptides of interest can also easily be sent back to the Galaxy History for further analysis – for example, using available tools for assessing functional impact of sequence variants identified via proteogenomics[18].

## Methods section

### Implementation

The MVP plugin is built on HTML5, CSS and JavaScript. The core of the MVP is based on standard JavaScript and open-source libraries. It receives data from a documented Galaxy SQLite data provider API. The main visualization is integrated into Galaxy via the Galaxy visualizations registry. Once registered, any dataset of type mz.sqlite will automatically be viewable from the MVP tool. The MVP tool uses the Data Tables[38] library to manage the presentation, sorting and filtering of data. It utilizes the Lorikeet MS/MS viewer[29, 39] to visualize PSMs and corresponding MS/MS spectra, and the IGV.js package[40] to interactively present features of interest, and within a single viewing window to visualize proteomic, transcriptomic and genomic data.

To install the MVP plugin within a Galaxy instance, the Galaxy visualization registry and plugin architecture is used. This component of Galaxy allow for custom visualizations of any recognized datatype. The MVP plugin is registered with Galaxy following the registry rules[28]. Every Galaxy instance has a visualization_plugins_directory available for custom visualizations. By default this directory is located in <your galaxy directory>/config/plugins/visualizations, but it can be set to any relative path. To install the MVP plugin, the .tar file is opened via the “untar” command and extracted in the visualization_plugins_directory. This creates a set of directories under mvpapp that contains all the code, css and HTML needed to run the plugin. The instance must be restarted to make the visualization accessible to the end-user.

### Accessibility

- The code and releases are available at: https://github.com/galaxyproteomics/mvpapplication-git.git
- The completed History used to demonstrate MVP functionality within the text above can be accessed by registering an account on the public Galaxy instance at usegalaxy.eu. Once logged in, go to Shared Data → Histories and search for “MVP_History”. Along with input files for analysis, this History contains all necessary output files to launch MVP and explore its functionalities and usage.

### Training and documentation

- We have made available documentation describing the use of MVP within a proteogenomics workflow within the online Galaxy Training Network resource. This documentation can be accessed at: https://training.galaxyproject.org/training-material/topics/proteomics/tutorials/proteogenomics-novel-peptide-analysis/tutorial.html
- Additionally, workflows and documentation are also available for the proteogenomic workflows which generate the data that ultimately acts as inputs to the MVP tool. Training documentation and information on access to these workflows is available as follows:

- Protein database generation from RNA-Seq data: https://training.galaxyproject.org/training-material/topics/proteomics/tutorials/proteogenomics-dbcreation/tutorial.html
- Sequence database searching for peptide identification: https://training.galaxyproject.org/training-material/topics/proteomics/tutorials/proteogenomics-dbsearch/tutorial.html

### Inputs

We have described in detail in the Operation section the inputs needed for full operation of MVP to view and explore multi-omic data. To summarize the inputs include:

- The data table in the mz.sqlite format which enables interactive queries of PSM information for efficient viewing and manipulation in MVP
- The MS/MS peak lists in standard MGF format, as well as the FASTA-formatted protein sequence database used for generating PSMs
- The variant_annotation table containing annotation of variant amino acid sequences compared to the reference genome and proteome.
- The table feature_cds_map provides a mapping of the expressed amino acid sequence for proteins identified from PSMs to each of the exons in the reference genome coding for the protein

The History made available on usegalaxy.eu (described above in the Accessibility section) contains all the input files necessary for full operation of MVP.

### Performance

The application’s performance, as perceived by the end-user, is dependent on the server infrastructure that Galaxy is hosted on, and the end-user’s local machine used in accessing the Galaxy web application. The MVP application relies on the existing Galaxy API framework. Therefore, the application will benefit from the existing Galaxy server infrastructure without any configuration needed from the application. API response from Galaxy to the MVP application will scale with the performance of the supporting server.

Though the underlying database (mzSQLite data type) is a simple SQLite3 database, care has been taken to optimize performance. During database construction, multiple indexes are generated for every table and each index is dedicated to an API call. Since the database is a read-only database, the overhead incurred from indexed based insertion is minimal. The minimal extra time needed to create multiple indexes is spent during the mzToSQLite tool run. No indexes are created, no insertions, updates or deletions occur while the MVP application is accessing the data.

The size of the underlying dataset is never known by the MVP application. Every SQL call for data is based on SQL LIMITS and OFFSETS no matter how small or large the mzSQLite database. Using the limited SQL data return, data tables page data to the user as the user scrolls through large datasets. Using this standard technique, we have run tests on ~6GB datasets. Even at this large of size, table scrolling performance is indistinguishable from datasets orders of magnitude smaller.

## Availability of supporting source code and requirements

**Project name:** MVP Application

**Project home page:** https://github.com/galaxyproteomics/mvpapplication-git.git

**Operating system(s):** Linux based systems for the Galaxy server. No restrictions for the end-user. Modern browsers are required.

**Programming language:** HTML, JS and CSS

**Other requirements:** https://github.com/galaxyproteomics/mzToSQLite. Installable from Bioconda as “mztosqlite”. Available as a Galaxy tool from https://github.com/galaxyproteomics/tools-galaxyp/tree/master/tools/mz_to_sqlite. SAM file format: https://samtools.github.io/hts-specs/SAMv1.pdf

**License:** MIT.

**RRID:** SCR_018077

## Additional Material

**Additional File 1** is included as a .pdf formatted file.

**Title:** Example visualization of novel splice junction peptide

**Description:** The additional file contains a figure showing visualization of a novel peptide containing both a single amino acid substitution and spanning a novel splice junction. Visualization both in the MVP Peptide-Protein Viewer is shown as well as the IGVjs viewer.

**Additional File 2** is included as a .pdf formatted file.

**Title:** Schema of databases and tables acting as input to MVP.

**Description:** The additional file contains Entity-Relationship diagrams of the schema for the three databases which act as input to MVP: the mzSQLite database, the variant_annotation table and the feature_cds_map table.

## Acknowledgements

The authors thank Mo Heydarian and Karen Reddy for the usage of data for developing and testing the described software. We also thank the Supercomputing Institute at the University of Minnesota for maintenance of hardware and software infrastructure used in this work. This work was funded in part by NIH/NCI grant U24CA199347 to Dr. Griffin and the Galaxy-P team.

The authors declare no competing interests with the work described here.

## References Cited

1. Mertins P, Mani DR, Ruggles KV, Gillette MA, Clauser KR, Wang P, et al. Proteogenomics connects somatic mutations to signalling in breast cancer. Nature. 2016;534 7605:55–62. doi:10.1038/nature18003.

2. Nesvizhskii AI. Proteogenomics: concepts, applications and computational strategies. Nat Methods. 2014;11 11:1114–25. doi:10.1038/nmeth.3144.

3. Polyakova A, Kuznetsova K and Moshkovskii S. Proteogenomics meets cancer immunology: mass spectrometric discovery and analysis of neoantigens. Expert Rev Proteomics. 2015;12 5:533–41. doi:10.1586/14789450.2015.1070100.

4. Ruggles KV, Krug K, Wang X, Clauser KR, Wang J, Payne SH, et al. Methods, Tools and Current Perspectives in Proteogenomics. Mol Cell Proteomics. 2017;16 6:959–81. doi:10.1074/mcp.MR117.000024.

5. Zhang B, Whiteaker JR, Hoofnagle AN, Baird GS, Rodland KD and Paulovich AG. Clinical potential of mass spectrometry-based proteogenomics. Nat Rev Clin Oncol. 2019;16 4:256–68. doi:10.1038/s41571-018-0135-7.

6. Eng JK, Searle BC, Clauser KR and Tabb DL. A face in the crowd: recognizing peptides through database search. Mol Cell Proteomics. 2011;10 11:R111 009522. doi:10.1074/mcp.R111.009522.

7. Armengaud J, Trapp J, Pible O, Geffard O, Chaumot A and Hartmann EM. Non-model organisms, a species endangered by proteogenomics. J Proteomics. 2014;105:5–18. doi:10.1016/j.jprot.2014.01.007.

8. Renuse S, Chaerkady R and Pandey A. Proteogenomics. Proteomics. 2011;11 4:620–30. doi:10.1002/pmic.201000615.

9. Jagtap PD, Johnson JE, Onsongo G, Sadler FW, Murray K, Wang Y, et al. Flexible and accessible workflows for improved proteogenomic analysis using the Galaxy framework. J Proteome Res. 2014;13 12:5898–908. doi:10.1021/pr500812t.

10. Afgan E, Baker D, Batut B, van den Beek M, Bouvier D, Cech M, et al. The Galaxy platform for accessible, reproducible and collaborative biomedical analyses: 2018 update. Nucleic Acids Res. 2018;46 W1:W537–W44. doi:10.1093/nar/gky379.

11. Boekel J, Chilton JM, Cooke IR, Horvatovich PL, Jagtap PD, Kall L, et al. Multi-omic data analysis using Galaxy. Nat Biotechnol. 2015;33 2:137–9. doi:10.1038/nbt.3134.

12. Chambers MC, Jagtap PD, Johnson JE, McGowan T, Kumar P, Onsongo G, et al. An Accessible Proteogenomics Informatics Resource for Cancer Researchers. Cancer Res. 2017;77 21:e43–e6. doi:10.1158/0008-5472.CAN-17-0331.

13. Guillot L, Delage L, Viari A, Vandenbrouck Y, Com E, Ritter A, et al. Peptimapper: proteogenomics workflow for the expert annotation of eukaryotic genomes. BMC Genomics. 2019;20 1:56. doi:10.1186/s12864-019-5431-9.

14. Maringer K, Yousuf A, Heesom KJ, Fan J, Lee D, Fernandez-Sesma A, et al. Proteomics informed by transcriptomics for characterising active transposable elements and genome annotation in Aedes aegypti. BMC Genomics. 2017;18 1:101. doi:10.1186/s12864-016-3432-5.

15. Verbruggen S, Ndah E, Van Criekinge W, Gessulat S, Kuster B, Wilhelm M, et al. PROTEOFORMER 2.0: Further Developments in the Ribosome Profiling-assisted Proteogenomic Hunt for New Proteoforms. Mol Cell Proteomics. 2019;18 8 suppl 1:S126–S40. doi:10.1074/mcp.RA118.001218.

16. galaxyp.org.

17. Sheynkman GM, Johnson JE, Jagtap PD, Shortreed MR, Onsongo G, Frey BL, et al. Using Galaxy-P to leverage RNA-Seq for the discovery of novel protein variations. BMC Genomics. 2014;15:703. doi:10.1186/1471-2164-15-703.

18. Sajulga R, Mehta S, Kumar P, Johnson JE, Guerrero CR, Ryan MC, et al. Bridging the Chromosome-centric and Biology/Disease-driven Human Proteome Projects: Accessible and Automated Tools for Interpreting the Biological and Pathological Impact of Protein Sequence Variants Detected via Proteogenomics. J Proteome Res. 2018;17 12:4329–36. doi:10.1021/acs.jproteome.8b00404.

19. Kumar P, Panigrahi P, Johnson J, Weber WJ, Mehta S, Sajulga R, et al. QuanTP: A Software Resource for Quantitative Proteo-Transcriptomic Comparative Data Analysis and Informatics. J Proteome Res. 2019;18 2:782–90. doi:10.1021/acs.jproteome.8b00727.

20. Kroll JE, da Silva VL, de Souza SJ and de Souza GA. A tool for integrating genetic and mass spectrometry-based peptide data: Proteogenomics Viewer: PV: A genome browser-like tool, which includes MS data visualization and peptide identification parameters. Bioessays. 2017;39 7 doi:10.1002/bies.201700015.

21. Li K, Vaudel M, Zhang B, Ren Y and Wen B. PDV: an integrative proteomics data viewer. Bioinformatics. 2019;35 7:1249–51. doi:10.1093/bioinformatics/bty770.

22. https://galaxyproject.org/develop/visualizations/#visualization-plugin-tutorial.

23. Vizcaíno JA, Mayer G, Perkins S, Barsnes H, Vaudel M, Perez-Riverol Y, et al. The mzIdentML Data Standard Version 1.2, Supporting Advances in Proteome Informatics. Molecular & Cellular Proteomics. 2017;16 7:1275–85. doi:10.1074/mcp.M117.068429.

24. Deutsch EW. File formats commonly used in mass spectrometry proteomics. Mol Cell Proteomics. 2012;11 12:1612–21. doi:10.1074/mcp.R112.019695.

25. https://genome.sph.umich.edu/wiki/SAM#What_is_a_CIGAR.3F.

26. Li H, Handsaker B, Wysoker A, Fennell T, Ruan J, Homer N, et al. The Sequence Alignment/Map format and SAMtools. Bioinformatics. 2009;25 16:2078–9. doi:10.1093/bioinformatics/btp352.

27. Ewing A and Semple C. Breaking point: the genesis and impact of structural variation in tumours. F1000Research. 2018;7 doi:10.12688/f1000research.16079.1.

28. https://galaxyproject.org/visualizations-registry/

29. https://github.com/UWPR/Lorikeet.

30. https://github.com/igvteam/igv.js/.

31. Menschaert G, Wang X, Jones AR, Ghali F, Fenyo D, Olexiouk V, et al. The proBAM and proBed standard formats: enabling a seamless integration of genomics and proteomics data. Genome Biol. 2018;19 1:12. doi:10.1186/s13059-017-1377-x.

32. Heydarian M, Luperchio TR, Cutler J, Mitchell CJ, Kim MS, Pandey A, et al. Prediction of Gene Activity in Early B Cell Development Based on an Integrative Multi-Omics Analysis. J Proteomics Bioinform. 2014;7 doi:10.4172/jpb.1000302.

33. Ross PL, Huang YN, Marchese JN, Williamson B, Parker K, Hattan S, et al. Multiplexed Protein Quantitation inSaccharomyces cerevisiaeUsing Amine-reactive Isobaric Tagging Reagents. Molecular & Cellular Proteomics. 2004;3 12:1154–69. doi:10.1074/mcp.M400129-MCP200.

34. Argentini A, Staes A, Gruning B, Mehta S, Easterly C, Griffin TJ, et al. Update on the moFF Algorithm for Label-Free Quantitative Proteomics. J Proteome Res. 2019;18 2:728–31. doi:10.1021/acs.jproteome.8b00708.

35. Millikin RJ, Solntsev SK, Shortreed MR and Smith LM. Ultrafast Peptide Label-Free Quantification with FlashLFQ. J Proteome Res. 2018;17 1:386–91. doi:10.1021/acs.jproteome.7b00608.

36. Gao J, Aksoy BA, Dogrusoz U, Dresdner G, Gross B, Sumer SO, et al. Integrative Analysis of Complex Cancer Genomics and Clinical Profiles Using the cBioPortal. Science Signaling. 2013;6 269:pl1–pl. doi:10.1126/scisignal.2004088.

37. Pratt D, Chen J, Welker D, Rivas R, Pillich R, Rynkov V, et al. NDEx, the Network Data Exchange. Cell Systems. 2015;1 4:302–5. doi:10.1016/j.cels.2015.10.001.

38. https://www.datatables.net/.

39. https://github.com/jmchilton/lorikeet.git.

40. https://github.com/igvteam/igv.js.git.

