## Additional File 1 for "Multi-omics Visualization Platform: An extensible Galaxy plug-in for multi-omics data visualization and exploration"

### Example visualization of novel splice junction peptide

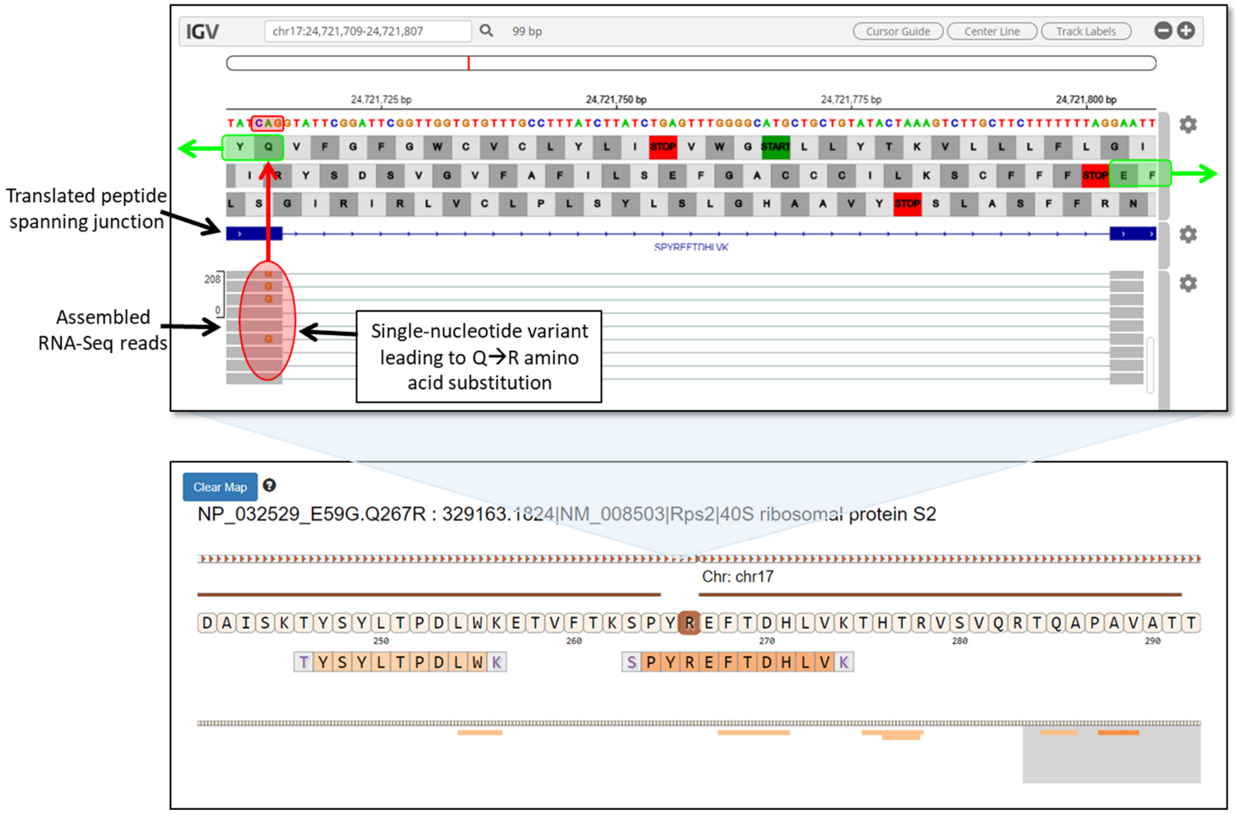


***Lower panel.*** The figure shows a potential novel peptide sequence identified and visualized in the Peptide-Protein Viewer. The peptide contains a single amino acid substitution at position 267, substituting an R (in dark shading) for the reference Q amino acid at this position.

***Upper panel.*** Opening up the IGV viewer from the Peptide-Protein Viewer shows that this peptide not only contains a single amino acid substitution, but also crossing a novel splice junction. The RNA-Seq reads indicated a G amino acid within the positive (sense) DNA strand (red shaded circle), indicating a codon sequence change from CAG to CGG in the genomic sequence, and leading to the substitution of R for Q at this position. The green shaded boxes show the amino acids within the identified peptide SP**YR-EF**TDHLVK which span a novel splice junction and are aligned with the RNA-Seq reads shown in gray in the lower track. The amino acids joined together across the junction and shown in the upper panel are in bold (--YREF--).
