## Additional File 2 for "Multi-omics Visualization Platform: An extensible Galaxy plug-in for multi-omics data visualization and exploration"

### Additional File 2-- Schema of databases and tables acting as input to MVP

#### 1) mzSQLite database schema

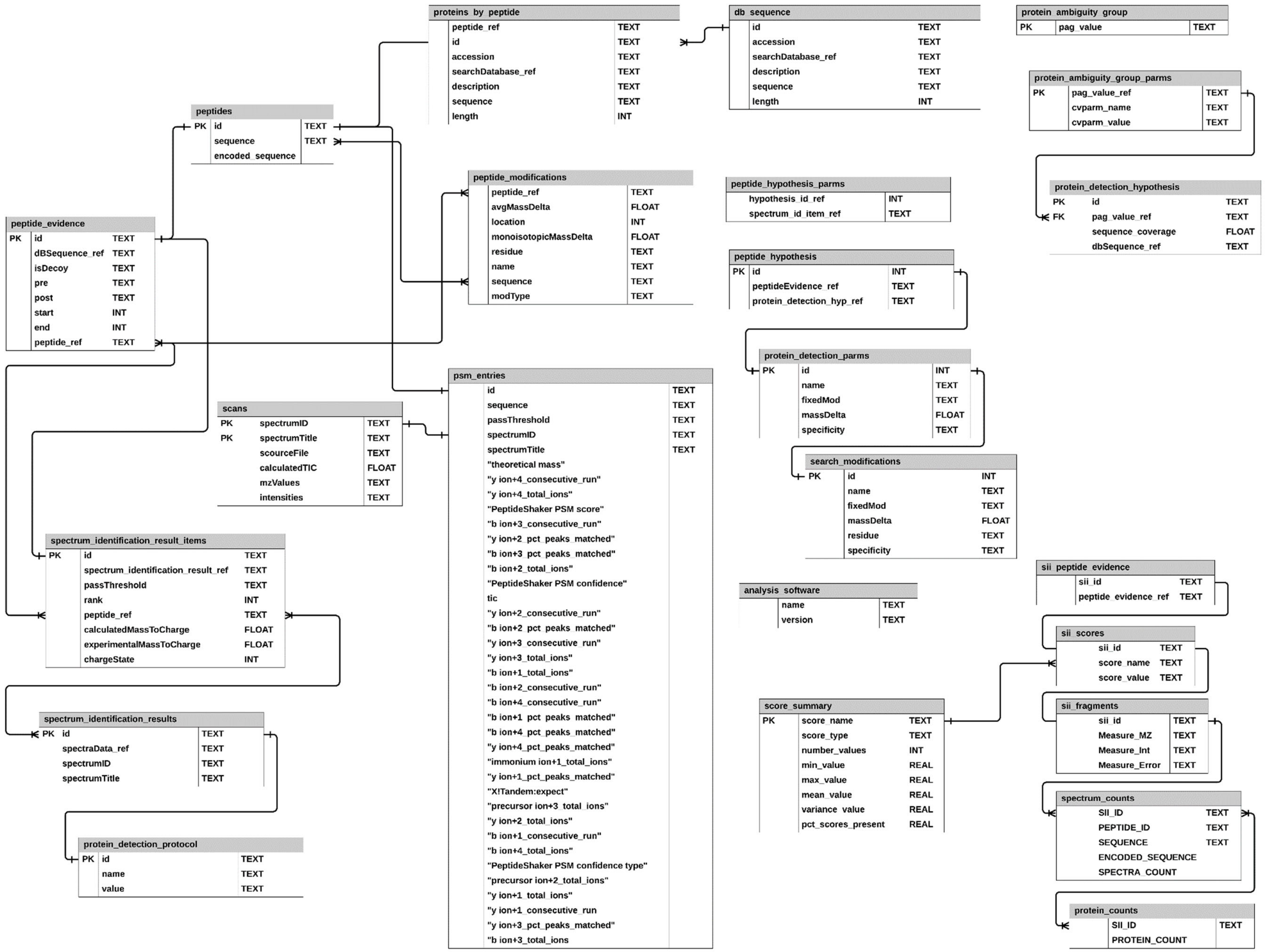

#### 2) variant_annotation table and the feature_cds_map table schema

| feature_cds_map | | |
| --- | --- | --- |
| | name | TEXT |
| | chrom | TEXT |
| | start | INT |
| | end | INT |
| | strand | TEXT |
| | cds_start | INT |
| | cds_end | INT |

| variant_annotation | | |
| --- | --- | --- |
| | name | TEXT |
| | reference | TEXT |
| | cigar | INT |
| | annotation | INT |
